# Combined efficacy of a novel antimicrobial cationic peptide polymer with conventional antibiotics to combat multi-drug resistant pathogens

**DOI:** 10.1101/735217

**Authors:** Kishore Reddy Thappeta, Yogesh Shankar Vikhe, Adeline Mei Hui Yong, Mary B. Chan Park, Kimberly A. Kline

## Abstract

Antibiotic-resistant infections are predicted to kill 10 million people worldwide per year by 2050 and to cost the global economy 100 trillion USD. Novel approaches and alternatives to conventional antibiotics are urgently required to combat antimicrobial resistance. We have synthesized a chitosan-based oligolysine antimicrobial peptide, CSM5-K5, which targets multidrug resistant (MDR) bacterial species. Here we show that CSM5-K5 exhibits rapid bactericidal activity against methicillin resistant *Staphylococcus aureus* (MRSA), MDR *Escherichia coli*, and vancomycin resistant *Enterococcus faecalis* (VRE). Combinatorial therapy of CSM5-K5 with antibiotics to which each organism is otherwise resistant restores sensitivity to the conventional antibiotic. CSM5-K5 alone significantly reduced pre-formed bacterial biofilm by two-four orders of magnitude and, in combination with traditional antibiotics, reduced pre-formed biofilm by more than two-three orders of magnitude at sub inhibitory concentrations. Moreover, using a mouse excisional wound infection model, CSM5-K5 treatment reduced bacterial burdens by one to three orders of magnitude, and acted synergistically with vancomycin and tetracycline to clear VRE and MDR *E. coli,* respectively. Importantly, little to no resistance against CSM5-K5 arose for any of the three MDR bacteria during 15 days of serial passage. This work demonstrates the feasibility and benefits of using this synthetic cationic peptide as alternative to, or in combination with, traditional antibiotics to treat infections caused by MDR bacteria.

## INTRODUCTION

Antibiotic resistance is a threat to global public health and sustainability (1). Antibiotic resistance currently accounts for an estimated 70,000 annual deaths globally, and in the absence of new therapeutics, infections caused by resistant “superbugs” could kill an additional 10 million people each year worldwide by 2050, surpassing cancer (2). Moreover, by 2050, antibiotic resistant infections are estimated to cost up to 3.5% of the global GDP, equivalent to 100 trillion USD (2). Because corporate antibiotic development pipelines of have progressively declined over the past 20 years (3), there is substantial interest in seeking alternative therapeutic approaches to combat these multidrug resistant (MDR) pathogens.

Host-derived antimicrobial peptides (CAMPs), typically composed of cationic and hydrophobic domains, have garnered interest as alternative therapies for MDR infections. CAMPs are electrostatically attracted to anionic bacterial cell surfaces, followed by peptide insertion into the lipid bilayer via their hydrophobic residues (4, 5). Chitosan is a polysaccharide composed of repeating N-glucosamine, with a structure similar to that of bacterial peptidoglycan. This unique feature of chitosan renders it potentially compatible with the bacterial cell wall. Numerous studies have examined the efficacy of chitosan-CAMP hybrids containing quaternary ammonium (6, 7), pyridinium (8), piperazinium (9), phosphonium (10), or sulfonamide (11) derivatives. However, many of these derivatives possess high cationicity and rely on hydrophobicity for improved bacterial interaction, which often lead to mammalian hemolysis and toxicity. To overcome these limitations, we have synthesized a low molecular weight copolymer CSM5-K5 hydrochloride salt (where CSM denotes chitosan monomer repeat units and K denotes lysine amino acid repeat units) with a controlled molecular weight of 1450 Da (12). CSM5-K5 displays low hemolysis and toxicity, is antibacterial against a variety of MDR bacteria, and reduces MRSA bacterial burden in a murine wound infection model by four orders of magnitude (12).

In this study, we examine the ability of CSM5-K5 to reduce pre-formed methicillin resistant *Staphylococcus aureus* (MRSA), MDR *Escherichia coli,* and vancomycin resistant *Enterococcus faecalis* (VRE) biofilm *in vitro* and *in vivo*. We show that CSM5-K5 alone and in synergy with clinically useful antibiotics demonstrates bactericidal activity and anti-biofilm activity both *in vitro* and *in vivo* in a mouse excision wound infection model. Taken together, these results demonstrate the feasibility and benefits of using the synthetic cationic peptide CSM5-K5 as alternative to, or in combination with, traditional antibiotics to treat difficult-to treat infections caused by MDR bacteria.

## MATERIALS AND METHODS

### Bacterial strains and growth conditions

Bacterial strains used in this study are listed in **Table 1**. *E. coli* strains were grown overnight in Luria-Bertani (LB) broth with shaking, or on agar at 37°C under static conditions. *E. faecalis* strains were grown statically in brain heart infusion (BHI) broth or agar at 37°C under static conditions. *S. aureus* strains were grown overnight in tryptic soy broth (TSB) or agar at 37°C under static conditions. We used Muller Hinton (MH) broth with shaking or with 1.5% agar to perform antibiotic susceptibility assays. All inoculations were cultured for 16-18 hours at 37°C, unless stated otherwise. Overnight cultures of bacteria were centrifuged at 6,000 *g* for 5 minutes and resuspended in 1× PBS at optical density (OD_600_) 0.7 for all MIC assays, unless stated otherwise.

**Table 1.**
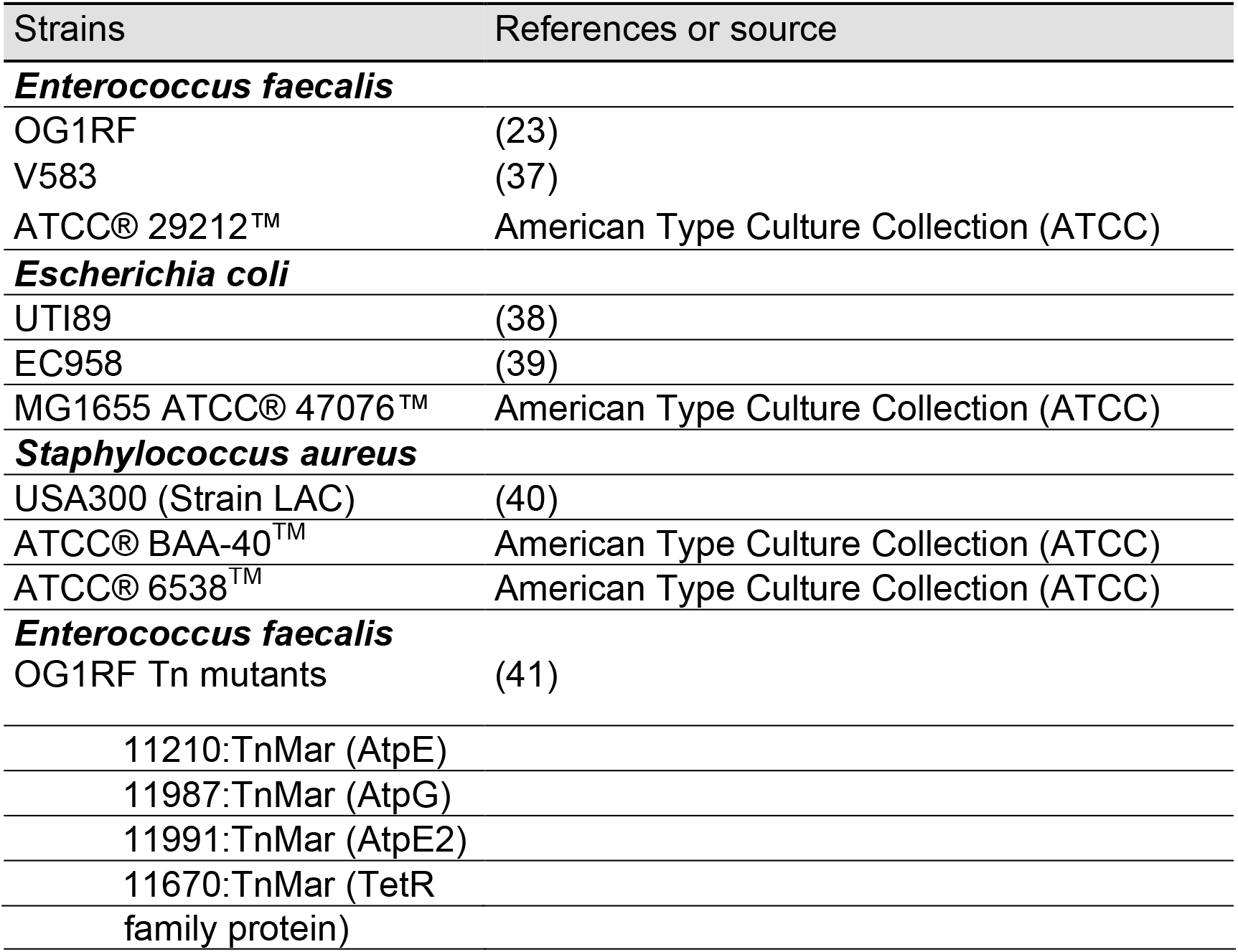
Bacterial strains used in this study.

### Polymer synthesis

CSM5-K5 was prepared as previously described in (12), with the following modifications. 5 g of low molecular weight chitosan (MW 200KDa) was first dispersed in 100 mL of anhydrous DMF and sonicated at 80°C for 1 hour under Argon protection. Protection of the amine group on chitosan was carried out by further adding 13.8 g of phthalic anhydride under 130°C and reacting for 24 hours. Further protection of 6-hydroxyl group was carried out by reacting 8 g phthalic protected chitosan with 24 g of trityl chloride in 100 mL of anhydrous pyridine at 100°C for 24 hours. Then, the chitosan macroinitiator was obtained by deprotection of phthalic group using hydrazine. Typically, 5 g of protected chitosan was deprotected by 100 mL of 50% hydrazine at 100°C for 24 hours. The protected chitosan-grafted-polylysine was synthesized by ring-opening polymerization of lysine N-carboxyanhydride (NCA) monomer initiated from the chitosan macroinitiator. Briefly, 1.32 g of lysine-NCA monomer was dissolved in 8 mL of anhydrous DMF and 112 mg of chitosan macroinitiator was dissolved in 3 mL of anhydrous DMF, and the chitosan macroinitiator solution was added into lysine-NCA monomer solution under Ar protection to initiate polymerization. The polymerization was carried out at room temperature for 3 days. Ultra-short CSM5-K5 cationic peptidopolysaccharide was obtained by acidic deprotection and hydrolysis of 1 g protected chitosan-grafted-polylysine with 10 mL of concentrated hydrochloride solution (37%) at 60°C for 100 mins. The crude deprotected product was neutralized by NaOH solution (1M) and dialyzed with 1000 Da cut-off cellulose membrane against deionized water for 5 days. The residue was lyophilized to obtain a white solid with molecular weight at 1450 Da (determined by MALDI-TOF analysis).

### Determination of minimum inhibitory concentration (MIC)

MIC values were determined using a broth micro dilution method as previously described (13). Bacterial cells were grown to mid-log phase and an optical density (600nm) of 0.5 for each organism, and normalized to 10^6^ CFU/mL. We dissolved the peptide polymer in water to a stock concentration of 10 mg/mL. The stock concentrations of antibiotics were prepared according to CLSI guidelines (14). Fifty microliters of the 1-5 x 10^5^ CFU/mL bacterial cultures were aliquoted into 96-well microtiter plates and mixed with 50 μL of media without or with two-fold dilutions of the peptide polymer or antibiotics and incubated for 16-18 hours at 37°C with shaking at 200 rpm. Growth inhibition was determined by measuring the optical density (OD_600_) of each well using a microplate reader (Infinite M200 Pro, Tecan, Switzerland). We determined the MIC of each bacterial strain by the lowest peptide concentration that inhibits more than 90% bacterial growth.

### Time-dependent killing assay

Bacteria were grown, diluted, and aliquoted into 96-well microtiter plates as described above, and mixed with 50 μL of either 0.5× or 1× MIC of the peptide polymer, with or without antibiotics added. The plates were sealed with parafilm and incubated at 37°C with shaking at 200 rpm. 20 μL of the culture was extracted at time intervals of 0, 0.5, 1, 2, 3, 5, and 24 hours for measurement of OD and colony-forming units, determined by dilution plating. 5 μL of each dilution was spotted on the BHI agar plates and incubated at 37°C for 24 hours prior to enumeration.

### Antimicrobial synergy assay

We measured the synergetic effects between the peptide polymer CSM5-K5 and antibiotics using the fractional inhibitory concentration index (FICI) method (15). We performed checkerboard susceptibility assays to measure MICs of antimicrobial combinations as previously described (16). The fractional inhibitory concentration (FIC) indices were calculated according to the following formulas:

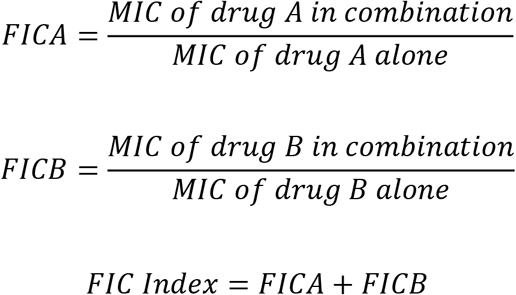

FIC index = FICA + FICB, where FICA = (MIC of drug A in combination)/ (MIC of drug A alone) and FICB = (MIC of drug B in combination)/ (MIC of drug B alone). Conservative interpretation of the FICI defines synergy as an FIC index of ≤ 0.5 (15).

### Minimum biofilm eradication concentration (MBEC) assay

MBEC biofilm assays were carried out according to published Innovotech methods (https://www.astm.org/Standards/E2799.htm?A). Briefly, overnight grown cultures of bacteria were diluted to between 10^5^ to 10^6^ CFU/mL in fresh MHB (for *E. faecalis* TSB with 0.25% glucose was used to achieve optimal growth), and 150 μL was transferred to the wells of a MBEC microtiter plate (Innovotech, Canada) and the MBEC lid was placed on top of the wells. Biofilms were grown on the MBEC pegs at 37°C with shaking at 200 rpm for 24 hours. The pegs were washed gently with 200 μL of PBS and the lid was transferred to a new plate in which wells contained of the CSM5-K5 and/or antibiotic in MHB and incubated at 37°C for 4 hours. The pegs were gently washed twice with 200 μL of PBS to remove non-adherent cells. Adherent biofilms on the pegs were placed in 200 μL of PBS and in a sonicating water bath for 30 min to disrupt the biofilm, prior to serial dilution in PBS CFU enumeration on BHI agar plates after growth at 37°C. Experiments were carried out in triplicate, and two independent experiments were executed for each of these assays.

### Antimicrobial resistance evolution assay

We asseessed resistance development of bacterial strains by sequential passaging each strain in the presence of sub-inhibitory concentrations of the cationic peptide polymer, CSM5-K5, or conventional antibiotics, essentially as previously described (17). In brief, bacterial cells were grown at 37°C in MH broth with shaking to mid-log phase and diluted to 1-5 × 10^5^ CFU/mL in MH broth containing 0.25×, 0.5×, 1×, 2× and 4× MIC concentrations of the peptide polymer or antibiotic and incubated at 37°C with shaking for EC958 and USA300 and statically for V583. At 24 hour intervals, the cultures from the second highest concentration of polymer or antibiotics that allowed growth (OD_600_ of 0.1-0.2) were diluted 1:100 into fresh media containing the same set of MIC concentrations, but based on the most recently visually observed MIC. The serial passaging was therefore repeated with increasing concentrations of peptide polymer or antibiotics over period of 15 days from two independent starter cultures per bacterial strain. To test stability of the resistant mutations, cultures that grew at the MIC or higher were streaked onto peptide polymer-free MH agar plates, individual colonies were selected and passaged daily in MH broth for five days, and a MIC was determined by broth micro dilution.

### Whole-genome sequencing

Genomic DNA was extracted from overnight bacteria cultures from resistant and wild type parental strains using the Wizard^®^ Genomic DNA Purification Kit (Promega, Madison, WI, USA), quantified and measured for DNA quality by Qubit High Sensitive dsDNA assay (Invitrogen, Carlsbad, CA, USA) and NanoDrop. The genomic DNA samples were sequenced on an Illumina MiSeq v3 platform. Whole genome sequencing data was analyzed using CLC Genomics Workbench 9.5 compared with reference genomes of *E. faecalis* V583, (Gene bank accession number NC_004668), *S. aureus* USA300 _FPR3757 (Genebank accession number NC_007793), and *E. coli* EC958 (Gene bank accession numberNZ_HG941718) from NCBI for mapping and annotation. Threshold variant frequency was set at >35% for all bacterial species. To detect variants, all mappings were analysed with the basic variant detection with regions of no coverage compared to the sequences of the respective wild type parental strains at the particular time point.

### Murine excisional wound model

The murine wound infection model was carried as described with minor modifications (18). Briefly, we grew the bacterial strains in 15 mL Tryptic Soy Broth supplemented with 0.25% glucose for 16-18 hours, at 37°C with continuous shaking at 200 rpm. Cells were collected, washed twice with 1 × sterile PBS and diluted to an OD0.5 and normalized to 1-3 x 10^5^ CFU/mL. We isoflurane-anesthetized groups of five male wild-type C57BL/6 mice (7-8 weeks old, 22 to 25 g; InVivos, Singapore) with their dorsal hair trimmed. Following trimming, Nair™ cream (Church and Dwight Co, Charles Ewing Boulevard, USA) was applied and the fine hair removed via shaving with a scalpel. We then disinfected the skin with 70% ethanol. A 6-mm biopsy punch (Integra Miltex, New York, USA) was used to create a full-thickness wound and 10^5^ CFU in 10 μL volume of the bacteria inoculum applied. We sealed the wound site with a Finn chamber on a Scanpor tape (Smart Practice, Phoenix, AZ, USA) and the chamber fixed to the skin via Fixomull stretch plasters (BSN medical GmbH, Hamburg, Germany). After 24 hours post infection, the Finn chambers were detached and discarded. We treated the infected wound with 10 μL of 1 ×MIC CSM5-K5 polymer prior to sealing of wound site with new Finn chambers and Scanpor tape and allow treatment for another 5 hours. After 5 hours of application, mice were euthanized and a one cm by one cm squared piece of skin surrounding the wound site was excised and collected in sterile 1 × PBS. We use a homogenizer (Pro200, SPD scientific, Singapore) for approximately 10 secs at high speed to homogenize the skin samples and the viable bacteria were enumerated by plating dilutions onto both BHI plates and antibiotic selection plates (ciprofloxacin for EC958, vancomycin for V583 and oxacillin for USA300) to ensure all recovered colony forming units corresponded to the inoculating strain. For synergy studies, we treated the infected wound with 5 μL of CSM5-K5 polymer and 5 μL of respective antibiotics. For each experiment, two independent biological replicates were performed containing 5 mice per group. Statistical analysis was performed by Mann-Whitney test using Prism software (GraphPad). We performed all approved procedures in accordance with the Institutional Animal Care and Use Committee (IACUC) in Nanyang Technological University, School of Biological Sciences (ARFSBS/NIEA0198Z) for murine wound infection model.

## RESULTS

### CSM5-K5 displays broad-spectrum killing against MDR bacterial strains

To extend our previous studies demonstrating CSM5-K5 (a cationic peptidopolysaccharide) displays broad-spectrum antimicrobial activity, we tested the efficacy of CSM5-K5 against a panel of MDR bacterial strains. Using minimum inhibitory concentration (MIC) assays, we found that CSM5-K5 displayed potent antibacterial activity against both Gram-positive and Gram-negative MDR pathogens including methicillin-resistant *S. aureus* USA300, vancomycin-resistant *E. faecalis* V583, and a highly virulent globally disseminating MDR and extended β-lactamase expressing strain of *E. coli* EC958. For all MDR strains tested, exposure to CSM5-K5 resulted in >98% killing within five hours (**Fig. 1A**).

**Figure 1:**
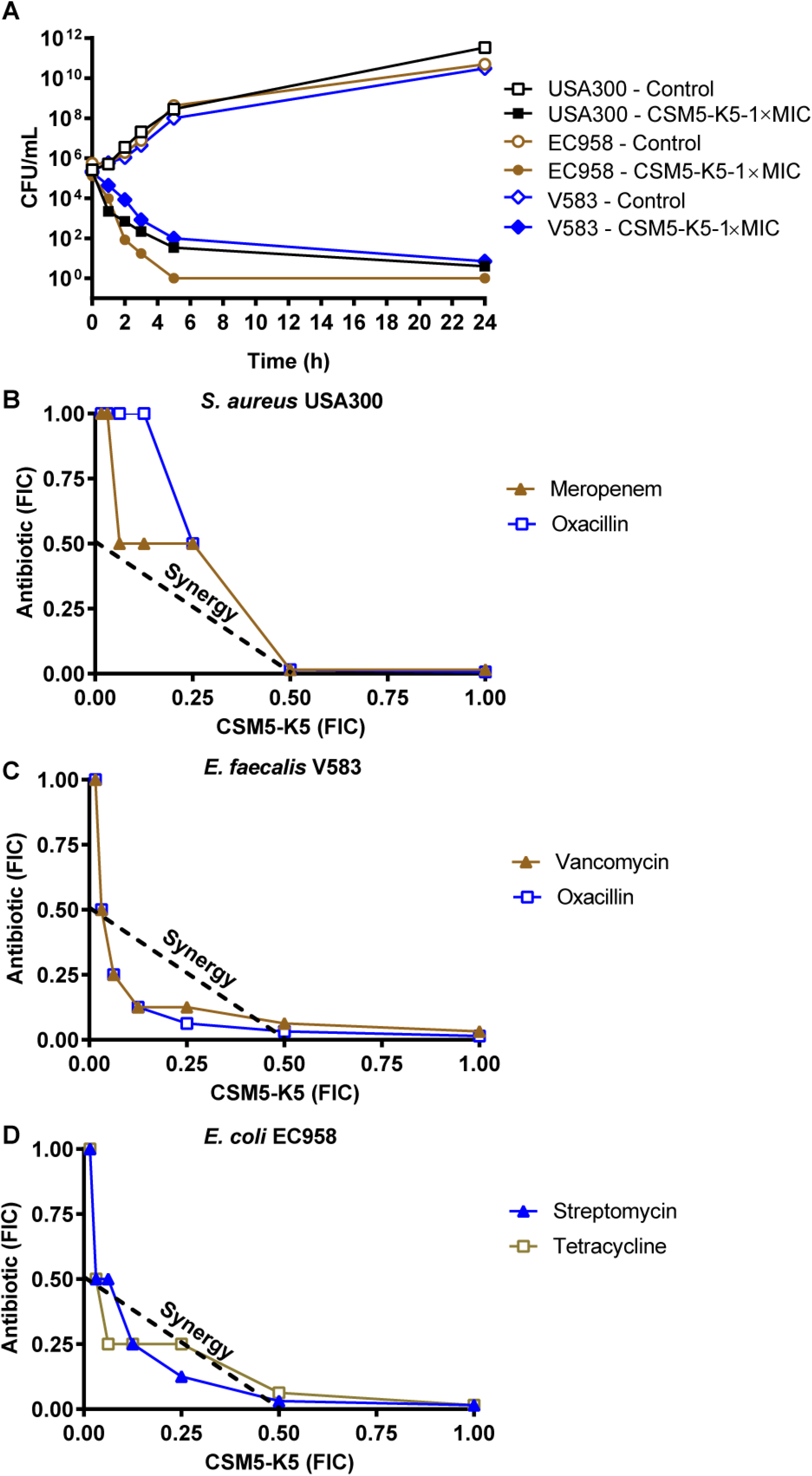
Synergistic combination treatment of MDR clinical isolates with CSM5-K5 and conventional antibiotics *in vitro*. (**A**) CSM5-K5 time-killing activity on the three MDR strains, *S. aureus* USA300, *E. coli* EC958 and *E. faecalis* V583. (**B-D**) The FIC of each antibiotic-CSM5-K5 pair for (**B**) *S. aureus* USA300 (**C**) *E. faecalis* V583 and (**D**) *E. coli* EC958 are plotted. Synergy is concluded when the sum of individual FIC values is <0.5 (indicated by diagonal dashed line). FIC values ≥0.5 indicates additive or partial synergistic activity. The data shown derive from two independent biological experiments.

### Combination treatment of CSM5-K5 with conventional antibiotics against MDR clinical isolates restores drug sensitivity

To determine whether the potency of CSM5-K5 could be further improved if used in combination with conventional antibiotics, we performed combinatorial MIC assays with clinically relevant antibiotics for MDR strains of *S. aureus* USA300, *E. faecalis* V583, and *E. coli* EC958. We first determined the MIC for CSM5-K5 and a panel of antibiotics against each MDR clinical isolate (**Table S1**) and then tested for synergy between CSM5-K5 and antibiotics which each organism was resistant to using a checkerboard assay containing two-fold dilutions for each compound. CSM5-K5 displayed partial synergy with oxacillin, meropenem, and other antibiotics against *S. aureus* USA300, with a fractional inhibitory concentration index (FICI) >0.5 to ≤0.1 (**Table S2 and Fig. 1B**). Further, reduction of CSM5-K5 to 0.5× MIC (8.0 μg/mL) in combination with oxacillin and meropenem reduced the respective MICs to 0.5 and 1.0 μg/mL, representing 64- and 8 fold reductions from their stand-alone MICs (32 and 8 μg/mL), respectively (**Table S1, data not shown**). Similarly, *E. faecalis* V583 is resistant to vancomycin and oxacillin, respectively, at concentrations of 32 μg/ml (**Table S1**). However, exposure of *E. faecalis* V583 to CSM5-K5 at 0.25× MIC (16 μg/mL) demonstrated synergistic activity with vancomycin and oxacillin, and restored drug sensitivity to MICs of 4 and 2 μg/mL, respectively (**Fig. 1C, data not shown**). Furthermore, CSM5-K5 exhibited potent synergy with the aminoglycosides streptomycin and tetracycline against *E. coli* EC958 (FICI <0.5) (**Fig. 1D, data not shown**). The MICs for *E. coli* EC958 for both streptomycin and tetracycline are 32 μg/mL (**Table S1**). However, CSM5-K5 (0.25× MIC) is synergistic with streptomycin and tetracycline at concentrations of ≤8 μg/mL, bringing the MICs to 8 and 4 μg/mL respectively (**data not shown**).

For all of the antibiotics we tested in combination with CSM5-K5, we observed reduction of the antibiotic MIC into the clinical therapeutic range for each bacterial species (14). We confirmed synergistic interactions using time killing curve assays and demonstrated that combinatorial bactericidal action occurred within synergistic concentrations of CSM5-K5 and respective antibiotics against each bacterial species. For *E. coli* EC958, we observed a more than two-log reduction within 5 hours of CSM5-K5 exposure at sub-MIC concentrations of 0.25× MIC and 0.37× MIC, with aminoglycosides streptomycin or tetracycline at 0.125× MIC or 0.25× (**Fig. S1A, B**). By contrast, higher concentrations of either individual antibiotic resulted in an initial drop in CFU followed by recovery by 5 hours and growth of the culture. Similarly, we observed a more than two-log reduction of *E. faecalis* V583 within 5 hours with combinatorial sub-MIC concentrations of CSM5-K5 and vancomycin or oxacillin (**Fig. S1C, D**). Moreover, we observed bactericidal action against *S. aureus* USA 300 at sub-MIC concentrations of CSM5-K5 of 0.5× with 0.03× MIC oxacillin and 0.06× MIC meropenem (**Fig. S1E, F**).

### CSM5-K5 alone and in combination with antibiotics effectively eradicates preformed biofilms of MDR pathogens *in vitro*

Many of the most difficult to treat nosocomial and chronic infections are biofilm-associated (19). The intrinsic antibiotic tolerance of biofilms coupled with genetic antibiotic resistance of MDR strains often renders these infections recalcitrant to treatment. Therefore, we next addressed whether CSM5-K5 alone and in combination with antibiotics had synergistic bactericidal activity against biofilm, similar to that observed for planktonic bacteria. We therefore performed a minimum biofilm eradication concentration (MBEC) assay in which we exposed pre-formed 27-28 hour biofilms to CSM5-K5 in the absence or presence of a conventional antibiotic for 3-4 hours. Exposure of *S. aureus* USA300, *E. faecalis* V583, or *E. coli* EC958 to 1 × MIC of CSM5-K5 resulted in an approximately three, two, or four-log reduction, respectively, for each of the organisms, representing >99% reduction in biofilm bacteria after just 4 hours of treatment compared to the untreated controls (**Fig. 2A**).

**Figure 2:**
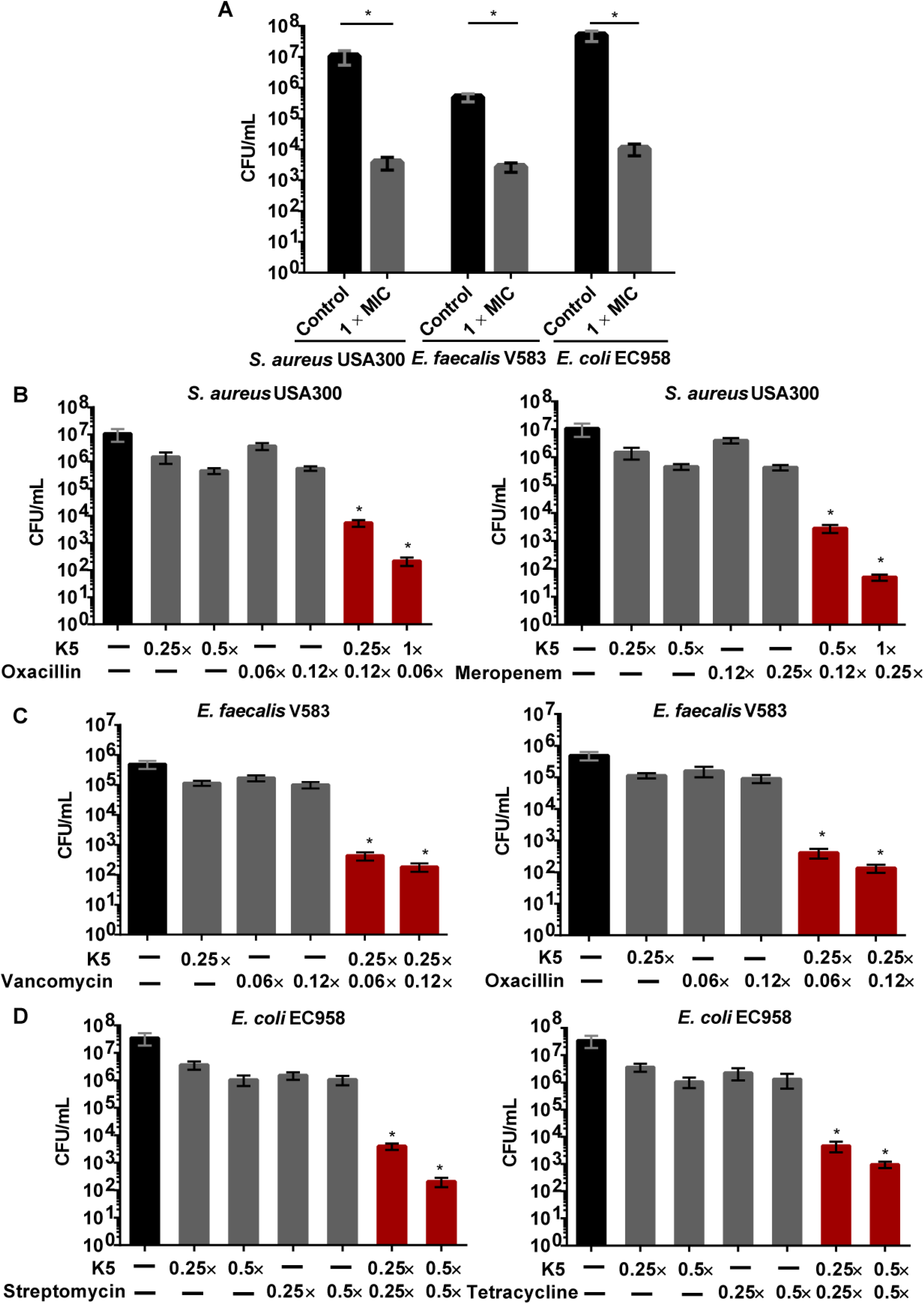
CSM5-K5 treatment reduces biomass of pre-formed MDR pathogen biofilms alone and synergistically in combination with traditional antibiotics *in vitro*. (**A**) *In vitro* biofilm assay upon exposure to CSM5-K5 (1 × MIC) for *S. aureus* USA300, *E. faecalis* V583, or *E. coli* EC958. *In vitro* biofilm assay upon exposure to sub-MIC concentrations of CSM5-K5 in combination with (**B**) oxacillin or meropenem against *S. aureus* USA300; (**C**) vancomycin or oxacillin against *E. faecalis* V583; (**D**) streptomycin or tetracycline against *E. coli* EC958. Data shown are combined from two independent experiments, each comprised of 3 biological replicates, with mean values ± standard error of mean plotted. Significant differences between groups analyzed by one-way ANOVA. (*P <0.05; **P ≤ 0.01; ***P ≤ 0.001; ****P ≤ 0.0001)

We observed for biofilms that, as for planktonic bacteria, CSM5-K5 was synergistic with conventional antibiotics at concentrations well below the MIC of each resistant strain. For example, CSM5-K5 displayed an antibiotic-enhancing effect for oxacillin and meropenem against *S. aureus* USA300 pre-formed biofilms at sub inhibitory concentrations of each. CSM5-K5 increased oxacillin biofilm reduction by 3-4.5 orders of magnitude compared with untreated biofilm, and 1.9-3.3 orders of magnitude compared to either of their single agent treatment alone (**Fig. 2B**). CSM5-K5 in combination with meropenem displayed an even higher efficacy for biofilm reduction, with 3.5-5.3 and 2-4 orders of magnitude reduction in CFU compared to untreated controls and single agent treatment, respectively (**Fig. 2B**). Similarly, we observed synergy of CSM5-K5 with vancomycin and oxacillin against *E. faecalis* V583 pre-formed biofilms of V583. Combinatorial treatment resulted in more than 3 and 2.4 log reductions compared to untreated controls and single agent treatment, respectively. At a sub-inhibitory concentration of CSM5-K5 at 0.25× MIC (16 μg/mL) with 0.06× MIC (2 μg/mL) and 0.12× MIC (4 μg/mL) of either of the two antibiotics, >99.8% of killing efficacy was achieved (**Fig. 2C**). Finally, we applied CSM5-K5 in combination with streptomycin and tetracycline to pre-formed *E. coli* EC958 biofilms. Combination of CSM5-K5 (at 0.25×-0.5× MIC) with either antibiotic at sub-inhibitory concentrations (0.25× and 0.5×) resulted in biofilm reduction of >2.5 - 4 and >3.8-5.2 orders of magnitude compared with single agent treated and untreated controls, respectively. Combination therapy killed >99% of *E. coli* EC958 biofilm (**Fig. 2D**).

### *In vivo* combined efficacy of CSM5-K5 with clinically relevant antibiotics

We next tested the ability of CSM5-K5 alone and in combination with conventional antibiotics to treat biofilm infections in a murine excisional wound infection model. Excisional wounds were infected with ~10^3^ CFU of each bacterial species, and then treated the wound site 24 hours post infection with CSM5-K5 at 1 × MIC alone or in combination with sub-inhibitory concentrations of antibiotics that each strain is resistant to. After 4-5 hours of antimicrobial treatment, CFU were enumerated from the infection site. Treatment of each bacterial species with 1 × MIC CSM5-K5 alone (16 μg/mL and 64 μg/mL for *S. aureus* USA300 and *E. faecalis* V583, respectively) resulted in greater than 90% reduction of USA300 and V583 CFU, compared to untreated controls (**Fig. 3A**). Treatment of *E. coli* EC958 with 1× MIC of CSM5-K5 resulted in >99% killing and a >3 log reduction in CFU compared to the untreated control (**Fig. 3A**). While combinatorial treatment did not further reduce *S. aureus* CFU at the concentrations tested in this experiment compared to CSM5-K5 alone (**Fig. 3A**), treatment of *E. faecalis* and *E. coli* infected wounds with sub-inhibitory concentrations of CSM5-K5 together with sub-inhibitory concentrations of antibiotics significantly improved bacterial killing compared to either single agent treated or untreated control (**Fig. 3C, D**). Importantly, combination treatment of each MDR strain with CSM5-K5 rendered them susceptible to antibiotic concentrations that fall below the clinical break point (0.06x MIC oxacillin = 2 μg/mL, 0.012 vancomycin= 4 μg/mL, 0.25x MIC streptomycin = 8 μg/mL). Hence, CSM5-K5 sensitized MDR strains of these wound-associated pathogens to antibiotics they are otherwise resistant.

**Figure 3:**
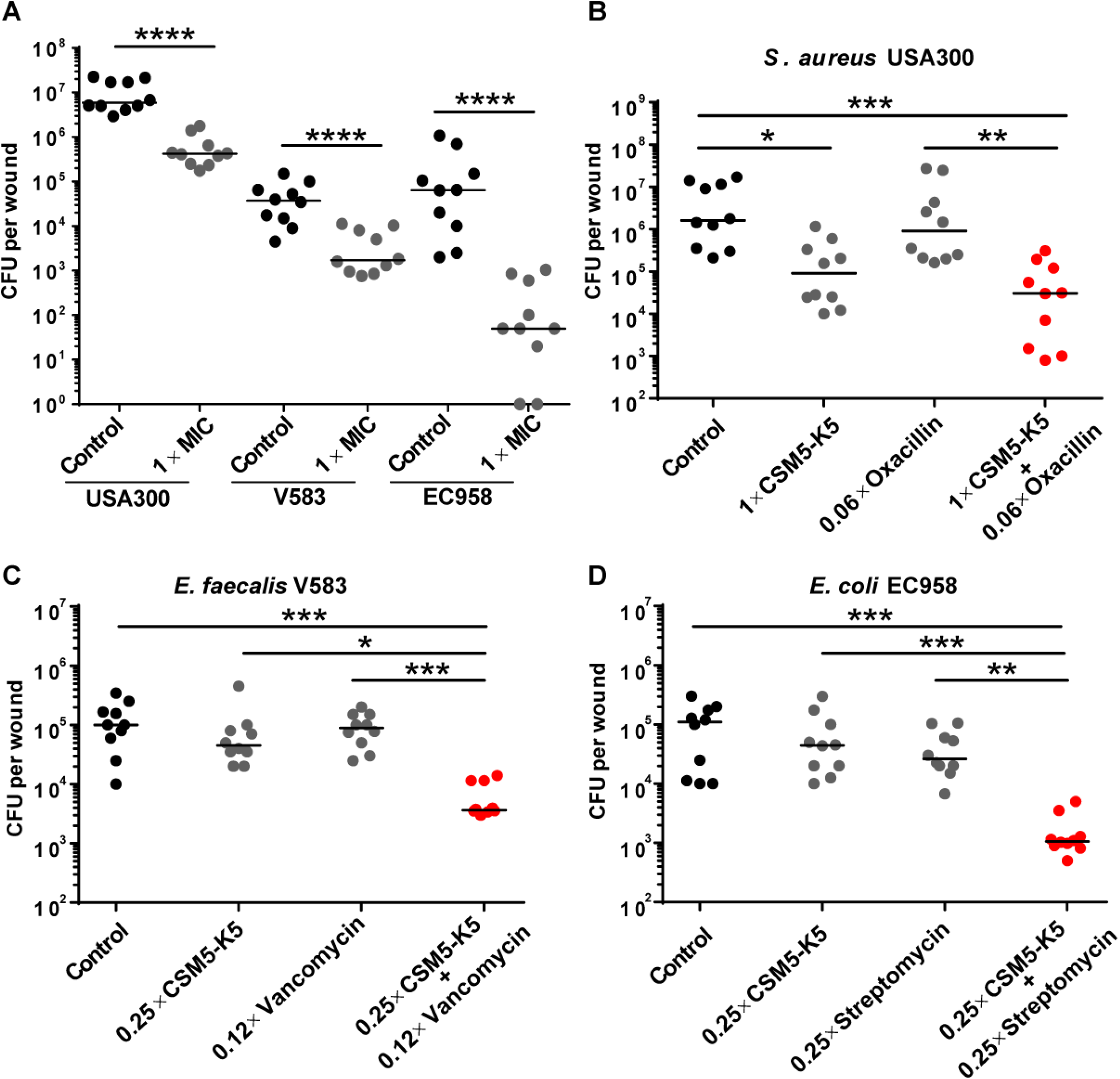
CSM5-K5 and conventional antibiotics synergistically attenuate MDR infection in a mouse model of biofilm-associated wound infection. Excisional wounds in mice were infected with the indicated pathogen for 24 hours followed by (**A**) treatment with CSM5-K5 (1× MIC) alone, or, in combination with antibiotics (**B-D**) for 5 hours. Each circle represents a single mouse. Horizontal lines represent the median of each group. Significant differences between groups were determined by the Kruskal-Wallis test (*P <0.05; **P ≤ 0.01; ***P ≤ 0.001; ****P ≤ 0.0001). Data are combined from 2 biological experiments, each containing 5 mice per group for each infection.

### CSM5-K5 exposure does not result in antimicrobial resistance

Development of antimicrobial resistance is defined by greater than a four-fold change to their initial MIC (20–23). To investigate if long-term use of CSM5-K5 can lead to resistance in *S. aureus* USA300, *E. faecalis* V583, and *E. coli* EC958, we subjected these three MDR strains to continuous serial passaging at sub-inhibitory concentrations of CSM5-K5 over 15 days. However, neither *S. aureus* USA300, *E. faecalis* V583, nor *E. coli* EC958 displayed more than a two-fold increase in MIC after 15 days of culture with sub-inhibitory concentrations of CSM5-K5 (**Fig. 4**). By contrast, serial passaging of *S. aureus* USA300 at sub-inhibitory concentrations of rifampicin, vancomycin, or gentamicin resulted in antibiotic resistance that was more than four-fold higher than the initial MIC (**Fig. 4A**). Similarly, MICs of rifampicin, tetracycline and meropenem were more than four-fold higher in *E. faecalis* V583 after 15 days of co-culture with the antibiotic (**Fig. 4B**). In *E. coli* EC958, resistance of 7, 6 and 3 times higher than the original MIC was observed for gentamicin, meropenem and polymyxin B, respectively (**Fig. 4C**). The failure to obtain CSM5-K5 resistance levels more than two-fold higher than their initial MIC values suggest a non-specific mode of action.

**Figure 4:**
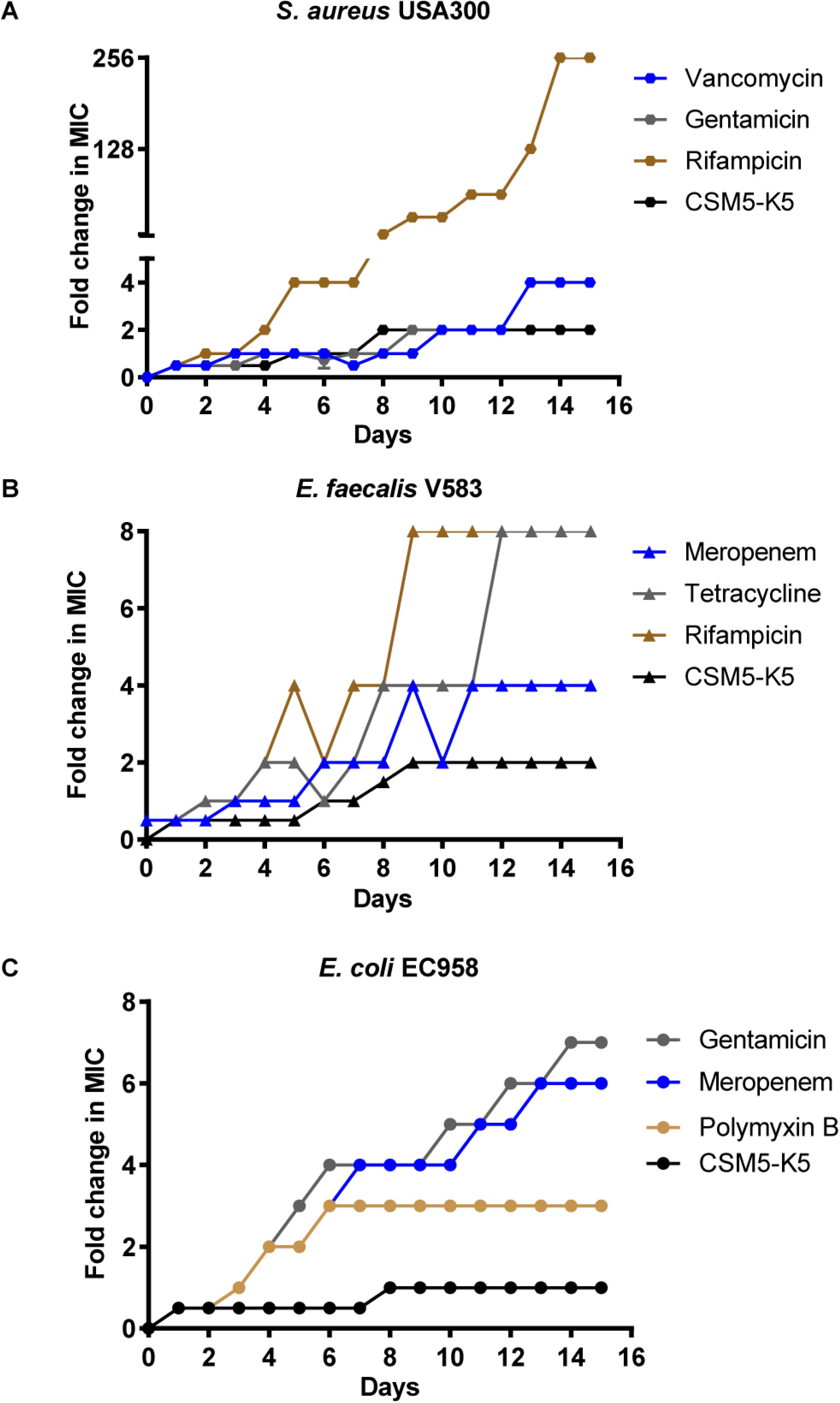
Prolonged CSM5-K5 exposure does not result in antimicrobial resistance. Continuous serial passaging of (**A**) *S. aureus* USA 300 (**B**), *E. faecalis* V583 (**C**) and *E. coli* EC958 at sub-inhibitory concentrations of CSM5-K5 or antibiotics over 15 days.

### Genetic basis of CSM5-K5^R^ in MDR *E. coli, S. aureus*, and *E. faecalis*

To identify the genetic basis for low level CSM5-K5^R^, we performed whole-genome sequencing (WGS) analysis of CSM5-K5^R^ isolates of *E. coli* EC958, *S. aureus* USA300 and *E. faecalis* V583. With the aim of detecting mutations related to CSM5-K5^R^ in *E. faecalis* V583, we sequenced 5 isolates from each of two independent serial passage experimental evolution assays (10 isolates in total) isolated at day 6 when the CSM5-K5 MIC was 64 μg/ml (1 × MIC). Using a threshold variant frequency cut-off of >35%, we found that resistant mutants contained a diverse set of mutations (**Table S3**). At day 6, we observed independent mutations in genes encoding ATP synthase machinery in 6 of 10 mutants (**Table S3, Fig. 4B**). We validated this finding by assessing the MIC for *E. faecalis* transposon mutants in the respective genes in strain OG1RF (*atpE, atpE2,* and *atpG*), which all displayed a 2-fold higher MIC for CSM5-K5 (**Table S4**). CSM5-K5^R^ *E. faecalis* mutants isolated at day 15 when the MIC reached 2x (128 μg/ml) appeared to be siblings and displayed mutations in *guaB,* and genes encoding a TetR family regulator and an HAD superfamily hydrolase in all isolates. Examination of a transposon mutant in the TetR family regulator, OG1RF_11670, which displays >99% sequence identity with V583 EF2066 demonstrated 2 fold higher MIC for CSMK5-K5. Together these results indicate that mutations in ATP synthase and a TetR family regulator can contribute to CSM5-K5 resistance in *E. faecalis*.

Similarly, we sequenced nine isolates of CSM5-K5^**R**^ *S. aureus* USA300 from two independent serial passage evolution experiments at day 5 at which MIC was 16 μg/mL (1× MIC) (**Table S3, Fig. 4A**). We identified a diverse array of mutations, including substitution mutations in the MFS transporter encoded by *narK* and in an ABC transporter permease (FtsX-like permease family protein), both of which are known to confer antimicrobial peptide resistance (24–26) and which may contribute to the observed low level CSM5-K5 resistance in *S. aureus*.

Nine CSM5-K5^R^ isolates of *E. coli* EC958 from two independent experiments were also isolated and sequenced at day 8, where MIC was 32 μg/mL (1× MIC) (**Table S3, Fig. 4C)**. All mutants contained a nonsense mutation in the membrane associated peptidase, encoded by *pepP,* which cleaves peptide bonds between any amino acid and proline (27) and is involved in outer membrane vesicle production (28), both of which could be contributing to the mechanism of CSM5-K5 resistance.

### Restored oxacillin susceptibility in CSM5-K5^R^ USA300 isolates

In an attempt to understand why we observed enhanced oxacillin susceptibility after *S. aureus* co-incubation with CSM5-K5 and oxacillin, we reanalyzed the same CSM5-K5^R^ mutants with a reduced threshold variant frequency of <35% and observed mutations in the *ebh* gene encoding hyperosmolarity resistance protein Ebh in nine isolates, which is associated with susceptibility to the β-lactam antibiotic oxacillin (29) (**Table S5**). We therefore examined whether *S. aureus* USA300 CSM5-K5^R^ isolates display oxacillin sensitivity and found that all *S. aureus* isolates showed reduced susceptibility to oxacillin (MIC ≤ 2 μg/mL) compared to wild type (MIC 32 μg/mL) (**Table S6**). *S. aureus* USA300 CSM5-K5^R^ isolates also displayed 2 to 4-fold greater susceptibility to carbenicillin and piperacillin, but no changed susceptibility to other antibiotic classes tested (**Table S6**). Similarly, CSM5-K5^R^ did not confer cross resistance to any other antibiotic for either *E. faecalis* or *E. coli* (**Table S6**). Finally, we selected and evaluated whether CSM5-K5^R^ mutant 11 conferred oxacillin susceptibility to this otherwise oxacillin resistant strain both *in vitro* and *in vivo* models. We observed that CSM5-K5^R^ mutant 11 exhibited oxacillin susceptibility at MIC 2 μg/mL (**Fig. 5A).** Similarly, we observed a significant difference between mutant and wild type growth when treated with oxacillin as well as with CSM5-K5 (**Fig. 5B).** Finally, in the murine excisional wound infection model, we observed that oxacillin alone was now effective in reducing infection by a CSM5-K5^R^ mutant 11 at 0.125× MIC (4 μg/mL) determined for this resistant strain (**Fig. 5C**).

**Figure 5:**
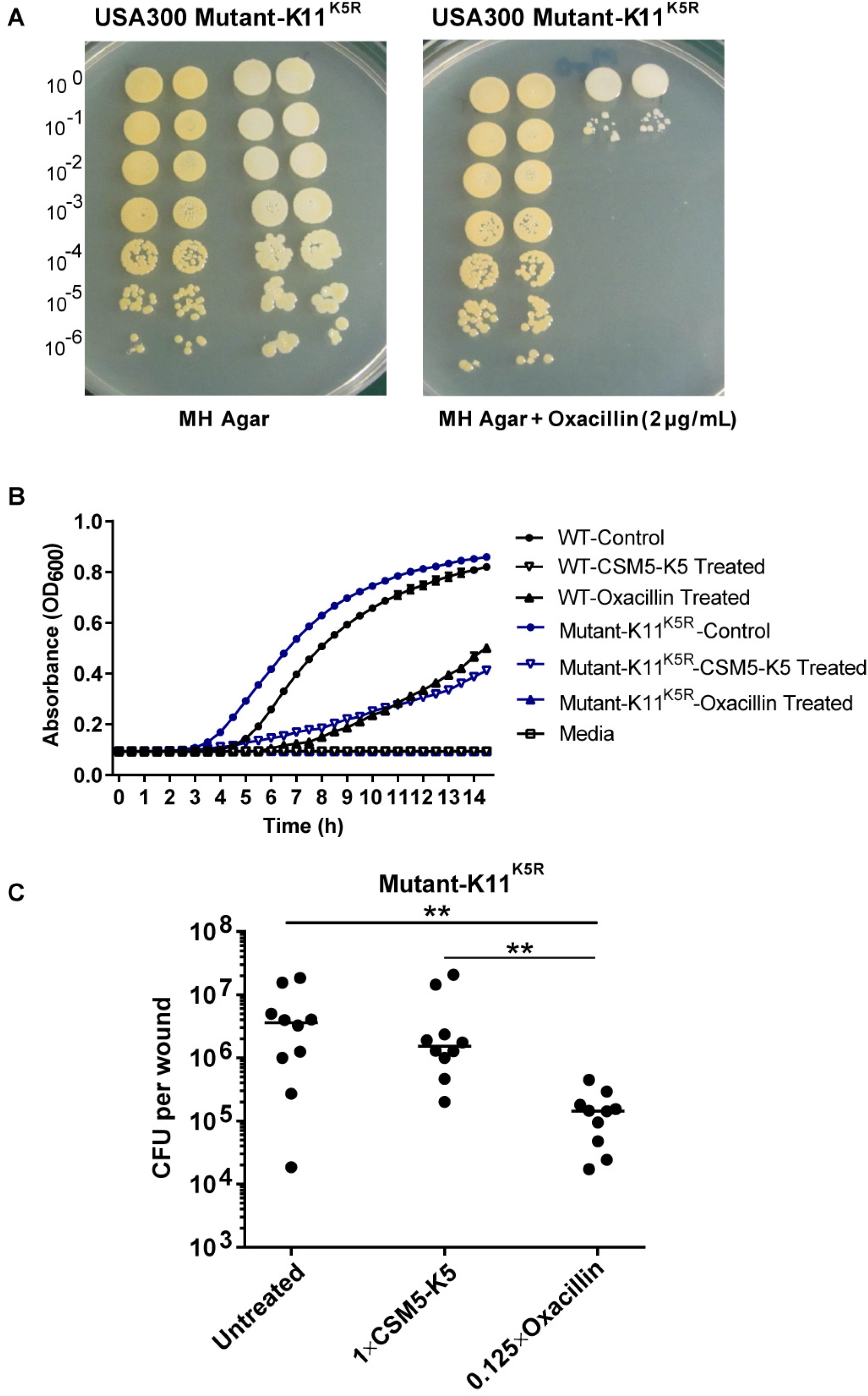
Collateral oxacillin sensitivity in *S. aureus* USA300 CSM5-K5^R^ mutants. (**A**) Serial dilutions of WT USA300 and CSM5-K5^R^ mutant-11 were spotted onto MH agar alone or containing 2 μg/mL oxacillin. (**B**) Growth curves of the *S. aureus* USA300 WT parent strain and CSM5-K5^R^ mutant-11 in the presence of oxacillin and CSM5-K5 at concentration of 2 μg/mL and 16 μg/mL, respectively. (**C**) Excisional wounds in mice were infected with CSM5-K5^R^ mutant-11for 24 hours followed by 5 hour treatment with CSM5-K5 16 μg/mL (1× MIC) or oxacillin (4 μg/mL). Each circle represents a single mouse. Horizontal lines represent the median of each group. Statistical analysis was performed using the Kruskal-Wallis test. Data shown are combined from two independent experiments, each containing 5 mice per group for each infection.

## Discussion

Multidrug resistant (MDR) bacterial infections are a growing and significant threat to public health, and are caused by species including methicillin resistant *S. aureus,* vancomycin resistant *E. faecalis,* and the globally disseminated CTX-M type ESBL-expressing strains of *E. coli* (30, 31). Antimicrobial peptides (AMPs) are of interest as alternatives to traditional antibiotics because of their broad spectrum antimicrobial properties and efficacy against MDR bacteria; however, to date they have met limited success due to their often non-selective toxicity (32–36). We previously reported that chitosan based cationic polymers showed antimicrobial activity against a variety of MDR pathogens (12, 20–22). In this study, we demonstrate that CSM5-K5 is synergistically active with traditional antibiotics against three antimicrobial resistant pathogens growing as biofilms *in vitro* and *in vivo*. Moreover, CSM5-K5 restores sensitivity of MRSA USA300 to oxacillin, *E. faecalis* V583 to vancomycin, and MDR and ESBL producing *E. coli* EC958 to streptomycin. Significantly, antibiotic sensitivity is restored to concentrations equal or below the clinical break point value for each antibiotic. Finally, prolonged exposure to CSM5-K5 did not give rise to clinically significant levels of resistance. Moreover, remarkably, the low level CSM5-K5 resistance to that did arise in *S. aureus* USA300 exhibited collateral sensitivity to β-lactam antibiotics including oxacillin, piperacillin and carbenicillin, restoring oxacillin and carbenicillin sensitivity to clinical breakpoint values. Together, these data present a viable combinatorial treatment strategy for difficult to treat antibiotic tolerant biofilm-associated infections, as well as those and genetically encoded antibiotic resistance, with minimal risk of antimicrobial resistance.

CSM5-K5 is a cationic nanoparticle that is self-assembled from chitosan-graft-oligolysine chains with ultralow molecular weight (1450 Da) that selectively kills bacteria with minimal toxicity toward mammalian cells (12). Hydrogen bonding within CSM5-K5 causes the polymer chains to aggregate into small nanoparticles to concentrate the cationic charge of the lysine. Upon contact with the bacterial membrane, these cationic nanoparticles synergistically cluster anionic membrane lipids and produce a greater membrane perturbation and antibacterial effect than would be achievable by the same quantity of charge if dispersed in individual copolymer molecules in solution (12). We observed partial of full antimicrobial synergy, against both Gram-negative and Gram-positive pathogens, between CSM5-K5 and nearly every antibiotic that we tested. We propose that the membrane-perturbing action of CSM5-K5 enables increased uptake and/or access of each antibiotic to its target. In this study, this synergy could be achieved at antibiotic concentrations to which the MDR pathogens were otherwise resistant. These observations suggest that this may be true for any antimicrobial resistant organism, regardless of the resistance mechanism or antibiotic mechanism of action. Together, the data in this manuscript present a viable combinatorial treatment strategy for difficult to treat antibiotic tolerant biofilm-associated infections, as well as those with genetically encoded resistance to traditional antibiotics, with minimal risk of antimicrobial resistance.

## ACKNOWLEDGMENTS

We are grateful to Jenny Dale and Gary Dunny for supplying us with *E. faecalis* OG1RF transposon mutants used in this study. This work was supported by the National Research Foundation and Ministry of Education Singapore under its Research Centre of Excellence Program, and by a Singapore Ministry of Education Tier 3 grant (MOE2013-T3-1-002).

---

**Supplementary Figure 1: Time-dependent killing analysis by CSM5-K5 in combination with conventional antibiotics**

Time-killing activity of (**A**) streptomycin (Strept) and (**B**) tetracycline (Tetra) against *E. coli* EC958; (**C**) vancomycin (Van) and (**D**) oxacillin (Oxa) against *E. faecalis* V583 and (**E**) oxacillin (Oxa) and (**F**) meropenem (Mero) against *S. aureus* USA300.

